# Repeats Of Unusual Size in Plant Mitochondrial Genomes: Identification, Incidence and Evolution

**DOI:** 10.1101/376020

**Authors:** Emily L. Wynn, Alan C. Christensen

**Affiliations:** School of Biological Sciences, University of Nebraska, Lincoln, Nebraska 68588-0666, USA

**Keywords:** Plant mitochondrial genomes, Repeated sequence, Genome rearrangement, Organelle genome evolution

## Abstract

Plant mitochondrial genomes have excessive size relative to coding capacity, a low mutation rate in genes and a high rearrangement rate. They also have non-tandem repeats in two size groups: a few large repeats which cause isomerization of the genome by recombination, and numerous repeats longer than 50bp, often found in exactly two copies per genome. It appears that repeats in the size range from several hundred to a few thousand base pair are underrepresented. The repeats are not well-conserved between species, and are infrequently annotated in mitochondrial sequence assemblies. Because they are much larger than expected by chance we call them Repeats Of Unusual Size (ROUS). The repeats consist of two functional classes, those that are involved in genome isomerization through frequent crossing over, and those for which crossovers are rare unless there are mutations in DNA repair genes, or the rate of double-strand breakage is increased. We systematically described and compared these repeats, which are important clues to mechanisms of DNA maintenance in mitochondria. We developed a tool to find non-tandem repeats and analyzed the complete mitochondrial sequences from 135 plant species. We observed an interesting difference between taxa: the repeats are larger and more frequent in the vascular plants. Analysis of closely related species also shows that plant mitochondrial genomes evolve in dramatic bursts of breakage and rejoining, complete with DNA sequence gain and loss, and the repeats are included in these events. We suggest an adaptive explanation for the existence of the repeats and their evolution.

## Introduction

It has long been known that plant mitochondrial genomes are much larger than those of animals (Ward, B. L. *et al*. 1981) and include significant amounts of non-coding DNA (Schuster, W. and A. Brennicke 1994). These genomes also often have repeats of several kb, leading to multiple isomeric forms of the genome (Folkerts, O. and M. R. Hanson 1989; Klein, M. *et al*. 1994; Palmer, J. D. and L. A. Herbon 1988; Palmer, J. D. and C. R. Shields 1984; Siculella, L. *et al*. 2001; Sloan, D. B. *et al*. 2010; Stern, D. B. and J. D. Palmer 1986). Plant mitochondrial genomes have very low mutation rates, but paradoxically have such high rearrangement rates that there is virtually no conservation of synteny (Drouin, G. *et al*. 2008; Palmer, J. D. and L. A. Herbon 1988; Richardson, A. O. *et al*. 2013; Wolfe, K. *et al*. 1987).

In addition to the large, frequently recombining repeats, there are often other repeated sequences in the size range of 1kb and lower (Arrieta-Montiel, M. P. *et al*. 2009; Forner, J. *et al*. 2005). Ectopic recombination between these homeologous repeats has been shown to increase when double-strand breakage is increased, or in plants mutant for DNA maintenance genes (Abdelnoor, R. V. *et al*. 2003; Shedge, V. *et al*. 2007; Wallet, C. *et al*. 2015). Understanding the repeats is critical to fully understanding the mechanisms of DNA maintenance and evolution in plant mitochondria, yet they have never been systematically identified and analyzed. In addition to being infrequently and inconsistently annotated and described in mitochondrial genome sequences, repeats are often described as long, short and intermediate-length (Arrieta-Montiel, M. P. *et al*. 2009; Davila, J. I. *et al*. 2011; Miller-Messmer, M. *et al*. 2012). The repeats are sometimes thought to be distributed into two size classes (one of up to several hundred bp and another of several kb), but this is derived from early studies of Arabidopsis and a few other species in which repeats were described and annotated.

The most likely hypothesis that explains the peculiar characteristics of plant mitochondrial genomes is that double-strand break repair (DSBR) is abundantly used in plant mitochondria, perhaps to the exclusion of nucleotide excision and mismatch repair pathways (Christensen, A. C. 2014; Christensen, A. C. 2018). Double-strand break repair is very accurate when the repair is template-based, accounting for the low mutation rate in genes, but the nonhomologous end-joining or break-induced-replication pathways can account for the creation of repeats and chimeric genes, expansions, and loss of synteny through rearrangements.

The lack of a coherent nomenclature and the inconsistent reporting and annotation of repeated sequences leads to a number of questions. What is the best way to discover and characterize them? Is the size distribution really bimodal in angiosperms? Are there repeats in the mitochondria of other groups of green plants? How do they differ between groups? And can they be followed through evolutionary lineages like genes? Are the repeats themselves somehow adaptive, or are they a side-effect of DSBR that is neutral or nearly neutral? The availability in recent years of complete mitochondrial genome sequences across a wide variety of taxa of green plants allows us to begin addressing these questions. We describe a computational strategy for finding non-tandem repeats within genomes. Using this tool we describe the phylogenetic distribution of repeats in both size classes, examine their evolution in a family of closely related angiosperms, and propose an hypothesis for the evolutionary significance of the repeats and the DSBR processes that produce them.

## Materials and Methods

### Sequence data and manipulation

DNA sequences were downloaded as FASTA format files from GenBank (https://www.ncbi.nlm.nih.gov/genbank/). BLAST searches (Altschul, S. F. *et al*. 1990) were done using version 2.7.1 on a Linux-based machine. In addition to the sequences shown in Table 1, mitochondrial genomes from several Brassica species were used to compare close relatives. These sequences are as follows: *Brassica carinata;* JF920287, *Brassica rapa;* JF920285, *Brassica oleracea fujiwase;* AP012988, *Brassica napus polima;* FR715249, *Brassica juncea;* JF920288. Alignments were done using the clustalW implementation in the VectorNTI 11.5 software package (ThermoFisher).

**Table 1.** List of species and mitochondrial DNA accession numbers

| Genus species | Group | Subgroup | Accession Number |
| --- | --- | --- | --- |
| *Auxenochlorella protothecoides* | Chlorophytes | Chlorophyta | KC843974.1 |
| *Botryococcus braunii* | Chlorophytes | Chlorophyta | LT545992.1 |
| *Bracteacoccus aerius* | Chlorophytes | Chlorophyta | KJ806265.1 |
| *Chlamydomonas eustigma* | Chlorophytes | Chlorophyta | BEGY01000520.1 |
| *Chlamydomonas reinhardtii* | Chlorophytes | Chlorophyta | EU306617.1 |
| *Chlorella* sp. ArM0029B | Chlorophytes | Chlorophyta | KF554428.1 |
| *Dunaliella salina* strain GN | Chlorophytes | Chlorophyta | KX641169.1 |
| *Eudorina* sp. | Chlorophytes | Chlorophyta | KY442294.1 |
| *Gloeotilopsis planctonica* | Chlorophytes | Chlorophyta | KX306823.1 |
| *Gloeotilopsis sarcinoidea* | Chlorophytes | Chlorophyta | KX306822.1 |
| *Hariotina* sp. MMOGRB0030F | Chlorophytes | Chlorophyta | KU145405.1 |
| *Kirchneriella aperta* | Chlorophytes | Chlorophyta | NC_024759.1 |
| *Lobosphaera incisa* | Chlorophytes | Chlorophyta | KP902678.1 |
| *Microspora stagnorum* | Chlorophytes | Chlorophyta | KF060942.1 |
| *Mychonastes homosphaera* | Chlorophytes | Chlorophyta | NC_024760.1 |
| *Neochloris aquatica* | Chlorophytes | Chlorophyta | NC_024761.1 |
| *Oltmannsiellopsis viridis* | Chlorophytes | Chlorophyta | DQ365900.1 |
| *Ostreococcus tauri* | Chlorophytes | Prasinophytes | CR954200.2 |
| *Ourococcus multisporus* | Chlorophytes | Chlorophyta | NC_024762.1 |
| *Polytoma uvella* | Chlorophytes | Chlorophyta | NC_026572.1 |
| *Prototheca wickerhamii* | Chlorophytes | Chlorophyta | U02970.1 |
| *Pseudendoclonium akinetum* | Chlorophytes | Chlorophyta | AY359242.1 |
| *Tetrademus obliquus* | Chlorophytes | Chlorophyta | CM007918.1 |
| *Trebouxiophyceae* sp. MX-AZ01 | Chlorophytes | Chlorophyta | JX315601.1 |
| *Ulva flexuosa* | Chlorophytes | Chlorophyta | KX455878.1 |
| *Ulva linza* | Chlorophytes | Chlorophyta | NC_029701.1 |
| *Nephroselmis olivacea* | Chlorophytes | Nephroselmidophyceae | AF110138.1 |
| *Bathycoccus prasinus* | Chlorophytes | Prasinophytes | NC_023273.1 |
| *Cymbomonas tetramitiiformis* | Chlorophytes | Prasinophytes | KX013548.1 |
| *Micromonas* sp. RCC299 | Chlorophytes | Prasinophytes | FJ859351.1 |
| *Monomastix* sp. OKE-1 | Chlorophytes | Prasinophytes | KF060939.1 |
| *Prasinoderma coloniale* | Chlorophytes | Prasinophytes | KF387569.1 |
| *Pycnococcus provasolii* | Chlorophytes | Prasinophytes | GQ497137.1 |
| *Pyramimonas parkeae* | Chlorophytes | Prasinophytes | KX013547.1 |
| *Megaceros aenigmaticus* | Anthocerotophyta | Anthocerotophyta | EU660574.1 |
| *Phaeoceros laevis* | Anthocerotophyta | Anthocerotophyta | GQ376531.1 |
| *Aneura pinguis* | Marchantiophyta | Marchantiophyta | KP728938.1 |
| *Marchantia polymorpha* | Marchantiophyta | Marchantiophyta | NC_001660.1 |
| *Pleurozia purpurea* | Marchantiophyta | Marchantiophyta | NC_013444.1 |
| *Treubia lacunosa* | Marchantiophyta | Marchantiophyta | JF973315.1 |
| *Anomodon attenuatus* | Bryophytes | Bryophytes | JX402749.1 |
| *Atrichum angustatum* | Bryophytes | Bryophytes | KC784956.1 |
| *Bartramia pomiformis* | Bryophytes | Bryophytes | KC784955.1 |
| *Brachythecium rivulare* | Bryophytes | Bryophytes | KR732319.1 |
| *Bucklandiella orthotrichacea* | Bryophytes | Bryophytes | KP742835.1 |
| *Buxbaumia aphylla* | Bryophytes | Bryophytes | KC784954.1 |
| *Climacium americanum* | Bryophytes | Bryophytes | KC784950.1 |
| *Codriophorus aciculare* | Bryophytes | Bryophytes | KP453983.1 |
| *Funaria hygrometrica* | Bryophytes | Bryophytes | KC784959.1 |
| *Hypnum imponens* | Bryophytes | Bryophytes | KC784951.1 |
| *Nyholmia gymnostoma* | Bryophytes | Bryophytes | KX578030.1 |
| *Orthotrichum callistomum* | Bryophytes | Bryophytes | KX578029.1 |
| *Oxystegus tenuirostris* | Bryophytes | Bryophytes | KT326816.1 |
| *Physcomitrella patens* | Bryophytes | Bryophytes | NC_007945.1 |
| *Ptychomnion cygnisetum* | Bryophytes | Bryophytes | KC784949.1 |
| *Sanionia uncinata* | Bryophytes | Bryophytes | KP984757.1 |
| *Sphagnum palustre* | Bryophytes | Bryophytes | KC784957.1 |
| *Stoneobryum bunyaense* | Bryophytes | Bryophytes | KX578031.1 |
| *Syntrichia filaris* | Bryophytes | Bryophytes | KP984758.1 |
| *Tetraphis pellucida* | Bryophytes | Bryophytes | KC784953.1 |
| *Tetraplodon fuegianus* | Bryophytes | Bryophytes | KT373818.1 |
| *Ulota phyllantha* | Bryophytes | Bryophytes | KX578033.1 |
| *Zygodon viridissimus* | Bryophytes | Bryophytes | KX711975.1 |
| *Chaetosphaeridium globosum* | Streptophyta | Charophyta | AF494279.1 |
| *Chara vulgaris* | Streptophyta | Charophyta | AY267353.1 |
| *Chlorokybus atmophyticus* | Streptophyta | Charophyta | NC_009630.1 |
| *Closterium baillyanum* | Streptophyta | Charophyta | NC_022860.1 |
| *Entransia fimbriata* | Streptophyta | Charophyta | KF060941.1 |
| *Klebsormidium flaccidum* | Streptophyta | Charophyta | KP165386.1 |
| *Nitella hyalina* | Streptophyta | Charophyta | JF810595.1 |
| *Roya obtusa* | Streptophyta | Charophyta | KF060943.1 |
| *Ophioglossum californicum* | Tracheophyta | Fern | NC_030900.1 |
| *Psilotum nudum* | Tracheophyta | Fern | NC_030952.1 |
| *Huperzia squarrosa* | Tracheophyta | Lycophyte | NC_017755.1 |
| *Selaginella moellendorffii* | Tracheophyta | Lycophyte | GL377739.1 |
| *Ginkgo biloba* | Tracheophyta | Gymnosperm | KM672373.1 |
| *Welwitschia mirabilis* | Tracheophyta | Gymnosperm | NC_029130.1 |
| *Cycas taitungensis* | Tracheophyta | Gymnosperm | AP009381.1 |
| *Aegilops speltoides* | Tracheophyta | Angiosperm | AP013107.1 |
| *Ajuga reptans* | Tracheophyta | Angiosperm | KF709392.1 |
| *Allium cepa* | Tracheophyta | Angiosperm | KU318712.1 |
| *Arabidopsis thaliana* | Tracheophyta | Angiosperm | BK010421 |
| *Batis maritima* | Tracheophyta | Angiosperm | KJ820684.1 |
| *Beta vulgaris* | Tracheophyta | Angiosperm | BA000024.1 |
| *Betula pendula* | Tracheophyta | Angiosperm | LT855379.1 |
| *Boea hygrometrica* | Tracheophyta | Angiosperm | JN107812.1 |
| *Brassica nigra* | Tracheophyta | Angiosperm | KP030753.1 |
| *Butomus umbellatus* | Tracheophyta | Angiosperm | KC208619.1 |
| *Cannabis sativa* | Tracheophyta | Angiosperm | KU310670.1 |
| *Capsicum annuum* | Tracheophyta | Angiosperm | KJ865410.1 |
| *Carica papaya* | Tracheophyta | Angiosperm | EU431224.1 |
| *Castilleja paramensis* | Tracheophyta | Angiosperm | KT959112.1 |
| *Citrullus lanatus* | Tracheophyta | Angiosperm | GQ856147.1 |
| *Cocos nucifera* | Tracheophyta | Angiosperm | KX028885.1 |
| *Corchorus capsularis* | Tracheophyta | Angiosperm | KT894204.1 |
| *Daucus carota subsp. sativus* | Tracheophyta | Angiosperm | JQ248574.1 |
| *Diplostephium hartwegii* | Tracheophyta | Angiosperm | KX063855.1 |
| *Eruca vesicaria* | Tracheophyta | Angiosperm | KF442616.1 |
| *Geranium maderense* | Tracheophyta | Angiosperm | NC_027000.1 |
| *Glycine max* | Tracheophyta | Angiosperm | JX463295.1 |
| *Gossypium hirsutum* | Tracheophyta | Angiosperm | NC_027406.1 |
| *Helianthus annuus* | Tracheophyta | Angiosperm | CM007908.1 |
| *Heuchera parviflora* | Tracheophyta | Angiosperm | KR559021.1 |
| *Hibiscus cannabinus* | Tracheophyta | Angiosperm | MF163174.1 |
| *Hordeum vulgare* | Tracheophyta | Angiosperm | AP017300.1 |
| *Hyoscyamus niger* | Tracheophyta | Angiosperm | KM207685.1 |
| *Ipomoea nil* | Tracheophyta | Angiosperm | AP017303.1 |
| *Lagerstroemia indica* | Tracheophyta | Angiosperm | KX641464.1 |
| *Liriodendron tulipifera* | Tracheophyta | Angiosperm | NC_021152.1 |
| *Lolium perenne* | Tracheophyta | Angiosperm | JX999996.1 |
| *Lotus japonicus* | Tracheophyta | Angiosperm | JN872551.2 |
| *Medicago truncatula* | Tracheophyta | Angiosperm | KT971339.1 |
| *Milletia pinnata* | Tracheophyta | Angiosperm | JN872550.1 |
| *Mimulus guttatus* | Tracheophyta | Angiosperm | JN098455.1 |
| *Nicotiana tabacum* | Tracheophyta | Angiosperm | BA000042.1 |
| *Oryza sativa Indica* | Tracheophyta | Angiosperm | NC_007886.1 |
| *Phoenix dactylifera* | Tracheophyta | Angiosperm | JN375330.1 |
| *Populus tremula* | Tracheophyta | Angiosperm | KT337313.1 |
| *Raphanus sativus* | Tracheophyta | Angiosperm | NC_018551.1 |
| *Rhazya stricta* | Tracheophyta | Angiosperm | KJ485850.1 |
| *Ricinus communis* | Tracheophyta | Angiosperm | HQ874649.1 |
| *Salix suchowensis* | Tracheophyta | Angiosperm | KU056812.1 |
| *Salvia miltiorrhiza* | Tracheophyta | Angiosperm | KF177345.1 |
| *Schrenkiella parvula* | Tracheophyta | Angiosperm | KT988071.2 |
| *Silene latifolia* | Tracheophyta | Angiosperm | HM562727.1 |
| *Sinapis arvensis* | Tracheophyta | Angiosperm | KM851044.1 |
| *Sorghum bicolor* | Tracheophyta | Angiosperm | DQ984518.1 |
| *Spinacia oleracea* | Tracheophyta | Angiosperm | KY768855.1 |
| *Tripsacum dactyloides* | Tracheophyta | Angiosperm | NC_008362.1 |
| *Triticum aestivum* | Tracheophyta | Angiosperm | AP008982.1 |
| *Vicia faba* | Tracheophyta | Angiosperm | KC189947.1 |
| *Vigna angularis* | Tracheophyta | Angiosperm | AP012599.1 |
| *Viscum album* | Tracheophyta | Angiosperm | NC_029039.1 |
| *Vitis vinifera complete* | Tracheophyta | Angiosperm | FM179380.1 |
| *Zea mays strain NB* | Tracheophyta | Angiosperm | AY506529.1 |
| *Ziziphus jujuba* | Tracheophyta | Angiosperm | KU187967.1 |

### Repeat Analysis

Custom Python scripts are in Supplementary Materials. The script ROUSFinder.py (Supplemental File S1) uses blastn to perform a pairwise ungapped comparison of a sequence with itself, both strands separately, using a word size of 50, E value of 10,000, reward for a match +1, penalty for a mismatch −9, percent identity cutoff 99%. The script then concatenates the two output files and the full length identity is deleted. Alignments are then sorted and compared to identify and remove duplicate repeats, and an output file of the repeats in fasta format is created. This output file is then used as a query with the genome as subject to locate every copy of that repeat, create a table, and a table of binned sizes. The output can also be formatted for GenBank annotation. MultipleRepeats.py (Supplemental File S2) automates running ROUSFinder.py on every sequence within a directory.

### Data Availability

The authors state that all data necessary for confirming the conclusions presented in this article are represented fully within the article, including python scripts in Supplemental Material and accession numbers of DNA sequences shown in Table 1. Supplemental Material available at FigShare.

## Results

### Repeats in plant mitochondrial genomes

The existence of large non-tandem repeats in plant mitochondrial genomes is well known by now, but they have not been systematically identified and analyzed. Prior studies used variations of BLAST (Altschul, S. F. *et al*. 1990) to find repeats (Alverson, A. J. *et al*. 2011a; Alverson, A. J. *et al*. 2010; Alverson, A. J. *et al*. 2011b; Liu, Y. *et al*. 2014) or REPuter (Hecht, J. *et al*. 2011; Kurtz, S. and C. Schleiermacher 1999). Other available software packages specifically identify tandem repeats, or repeats matching known repetitive sequences. Due to the ready availability of BLAST and the flexibility of its use, and because most prior work used it, we wrote and used a Python script called ROUSFinder.py that uses BLAST to identify non-tandem repeats within mitochondrial genomes. The parameters for identification of a sequence repeat were relatively stringent and included a blastn word size of 50, a percent identify cutoff of 99% and match/mismatch scores of +1/-9. Any choice of parameters will necessarily identify some false positives and false negatives. These parameters were chosen in order to find duplicate copies of sequence that were either recently created or recently corrected by gene conversion. A duplication longer than 100 bases that has several mismatches in the center of the repeat unit will be identified as two different repeats in this way. However, the mismatches in the center are indicative of either two independent events producing the two parts of the repeat, or mutation and drift that have escaped gene conversion. Because we are concerned with the recombination behavior of the repeats we therefore choose to call these two different repeats. To analyze and identify repeats in a single sequence for further study or annotation would require additional manual curation of the output.

The species we used represent a significant subset of the complete mitochondrial genome sequences from green plants in GenBank and are shown in Table 1. Sequences available on GenBank are not a random sample across taxa (food crops are very over-represented, for example), so to reduce sampling bias somewhat we used only one species per genus. Incomplete sequences or sequences with gaps are not handled well by BLAST without further curation, so these were not used. Species with multiple distinct chromosomes were also not used because of the additional layer of complexity from inter- and intra-chromosomal repeats. The full output is in Supplemental Table S1. The repeats seen in plant mitochondrial genomes are much larger than those found in random sequence (data not shown), suggesting that they arise from specific biological processes and are not stochastic. For this reason we call them “Repeats Of Unusual Size” or ROUS (Christensen, A. C. 2018).

BLAST is an excellent tool for identification of repeated sequences. Our script automates the task of identifying repeats in both direct and inverted orientations, removes the full-length match, and provides the output in a convenient format that can be used for annotation or in spreadsheets for further analysis.

### Phylogenetic clustering

The distribution of repeat sizes forms distinct clusters between the phylogenetic groups (see Figure 1). Because there are different numbers of species in each group, and some species have an order of magnitude more total repeats than others, we represent the data as the fraction of species within that group that have at least one repeat within a given size range. The complete output is in Supplemental Table S1. There are several complete mitochondrial genomes from chlorophytes and bryophytes to compare to angiosperms. Within the chlorophytes, repeats of greater than 200bp are rare. The exceptions are the prasinophytes (discussed below) and a few interesting cases. *Chlamydomonas reinhardtii* has a 532 bp inverted repeat at the termini of its linear chromosome. *Dunaliella salina, Kirchneriella aperta* and *Polytoma uvella* have novel structures at a small number of loci that consist of overlapping and nested repeats and palindromes (Smith, D. R. *et al*. 2010). The function of these structures is unknown, but they are unusual and not common in the chlorophytes. The prasinophyte group resembles the rest of the chlorophytes in having no ROUS greater than 200bp but many of them include two copies of a single large repeat between 9.5 and 14.4 kb. This is similar to many chloroplast genomes and it is possible that this structure is involved in replication (Bendich, A. J. 2004). The bryophytes generally resemble the chlorophytes; there are no ROUS greater than 200bp.

**Figure 1.**
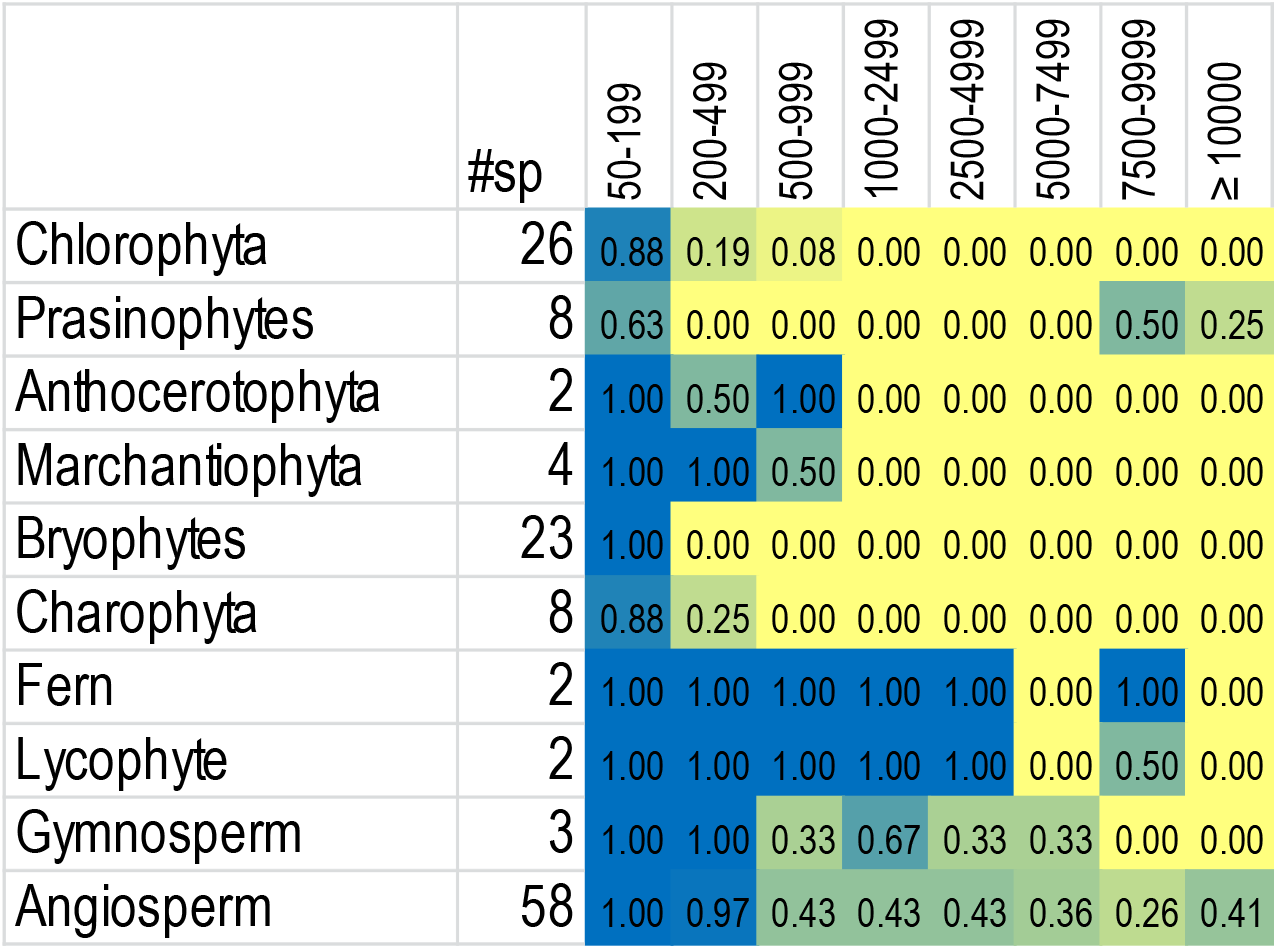
Size distributions of repeats in groups of species. The number of species represented in each group is shown. Headings indicate the bins of repeat sizes and the numbers indicate the fraction of species in that group that have at least one repeat of that size. Heat map color coding is blue for the highest value and yellow for zero.

In contrast to the chlorophytes and bryophytes, the Marchantiophyta (liverworts) and Anthocerotophyta (hornworts) have ROUS greater than 200bp in size, but none bigger than 867bp. Few taxa within the liverworts and hornworts have been sequenced, so this group is underrepresented, but the members available to date are consistent. The other lineages of streptophytic green algae (referred to as charophytes in GenBank) resemble the chlorophytes albeit with a slightly higher upper limit. In this group the largest repeat is found in *Chlorokybus atmophyticus* and is 291bp.

The ferns and lycophytes are strikingly different from the previous groups. Unfortunately the number of species sequenced is low. They have large numbers of repeats and the repeat sizes range well above 200bp, up to 10 kb. Some members of these groups, such as Huperzia, are similar to the bryophytes, but others are large and have significant repeat content (Guo, W. *et al*. 2017). These groups are also very underrepresented among available mitochondrial sequences (in part due to the complexity caused by the repetitive nature of the genomes (Grewe, F. *et al*. 2009)), but the patterns are noticeably different from the nonvascular plants described above.

The angiosperms are represented very well in the sequence databases. Only one member of this group does not have any ROUS above 200bp *(Medicago truncatula)*. A small number of angiosperms, scattered among plant families, lack repeats larger than 1 kbp, and approximately half include repeats larger than 9 kbp. *Silene conica*, a species with multiple large chromosomes not included in our dataset has a nearly 75kb sequence found in both chromosomes 11 and 12 (Sloan, D. B. *et al*. 2012). Gymnosperms are also underrepresented, but appear to be similar to the other vascular plants. Interestingly, the gymnosperms *Ginkgo biloba* and *Welwitschia mirabilis* resemble angiosperms, while *Cycas taitungensis* is more similar to ferns. The *C. taitungensis* mitochondrion has numerous ROUS, including many that are tandemly repeated. Five percent of this genome consists of the mobile Bpu element, a remarkable level of repetitiveness (Chaw, S. M. *et al*. 2008).

It is only in the vascular plants that the number and size of repeated sequences in mitochondrial genomes has been expanded. The vascular plants generally only have genomes a few times larger than the bryophytes, liverworts and hornworts, but the repeats are expanded well beyond proportionality to size. In addition, random sequences of comparable length do not have any repeats of the sizes discussed here (data not shown). Some taxa, such as the Geraniaceae, Plantago, and Silene include species with significantly expanded genomes (Park, S. *et al*. 2015; Parkinson, C. L. *et al*. 2005; Sloan, D. B. *et al*. 2012). These species are outliers in the magnitude of the genome sizes and number of repeats, but the underlying processes are likely to be the same. The overall picture is that there was a significant change in mitochondrial DNA maintenance mechanisms roughly coincident with the origin of the vascular plants.

### Repeat sizes and frequency in angiosperms

Large repeats of several kilobases have been identified in several species and shown to be recombinationally active, isomerizing angiosperm mitochondrial genomes (Folkerts, O. and M. R. Hanson 1989; Klein, M. *et al*. 1994; Palmer, J. D. and L. A. Herbon 1988; Palmer, J. D. and C. R. Shields 1984; Siculella, L. *et al*. 2001; Sloan, D. B. 2013; Stern, D. B. and J. D. Palmer 1986). A few species have been reported to lack such structures (Palmer, J. D. 1988). The first comprehensive catalog of repeated sequences shorter than 1000 base pairs was done in *Arabidopsis thaliana*, and they were shown to be recombinationally active in some mutant backgrounds, but not generally in wild type (Arrieta-Montiel, M. P. *et al*. 2009; Davila, J. I. *et al*. 2011; Miller-Messmer, M. *et al*. 2012; Shedge, V. *et al*. 2007). Is the spectrum of repeat sizes in Arabidopsis, and its bimodality, typical for angiosperms? Figure 2 illustrates the presence of repeats in the size range of 50bp to over 10,000 bp in 58 angiosperms. The overall pattern is that there is a multimodal distribution of sizes that is often bimodal. Gaps in the distribution are indicated in yellow in Figure 2, however, the size cutoffs of the gap are somewhat variable. Most species have a paucity of repeats between 600 and 10,000bp. Twelve of the 58 species have no repeats larger than 600bp, leaving open the question of if they isomerize through recombination or not. All of the other species have a large repeat of somewhere between 800bp and 65kbp. The total length of repeats in a species does not correlate with genome size (linear regression r^2^ = 0.08, data not shown), additional evidence that these are not produced by stochastic processes, and suggesting that they occur and change faster than speciation does.

### Alignment of repeats within the Brassicales

Understanding the evolution of the repeated sequences requires analysis of homologous repeats in related species. Of the species with sequenced mitochondrial genomes, the Brassicales order of plants has a number of such species. Within the *Brassica* genus there are three diploid species: *Brassica rapa, Brassica nigra* and *Brassica oleracea*, and three allotetraploid species (Cheng, F. *et al*. 2017). The diploid nuclear genomes are called the A, B and C genomes, respectively. Based on both nuclear and mitochondrial sequences it appears that *Brassica carinata* has the *B. nigra* and *B. oleracea* nuclear genomes (BBCC) and the *B. nigra* mitochondrial genome, while *Brassica juncea* has the *B. nigra* and *B. rapa* nuclear genomes (BBCC) and the *B. rapa* mitochondrial genome. *Brassica napus* is of two subspecies, *polima* and *napus*. Both have the *B. oleracea* and *B. rapa* nuclear genomes (AACC), but *B. napus polima* appears to have the *B. rapa* mitochondrial genome and *B. napus napus* has the *B. oleracea* mitochondrial genome (Chang, S. *et al*. 2011; Franzke, A. *et al*. 2011; Grewe, F. *et al*. 2014). Thus it appears that the hybridization event between *B. oleracea* and *B. rapa* occurred at least twice, with each species being the maternal parent. In the analysis below we use the *B. napus polima* mitochondrial genome. We compared these Brassica species to *Raphanus sativus* and *Sinapis arvensis*. These species are the closest relatives of the Brassicas with complete mitochondrial genome sequences (Grewe, F. *et al*. 2014). Several of these species were mapped prior to genomic sequencing, and repeated sequences and isomerization of the genomes was observed (Palmer, J. D. 1988; Palmer, J. D. and L. A. Herbon 1986).

**Figure 2.**
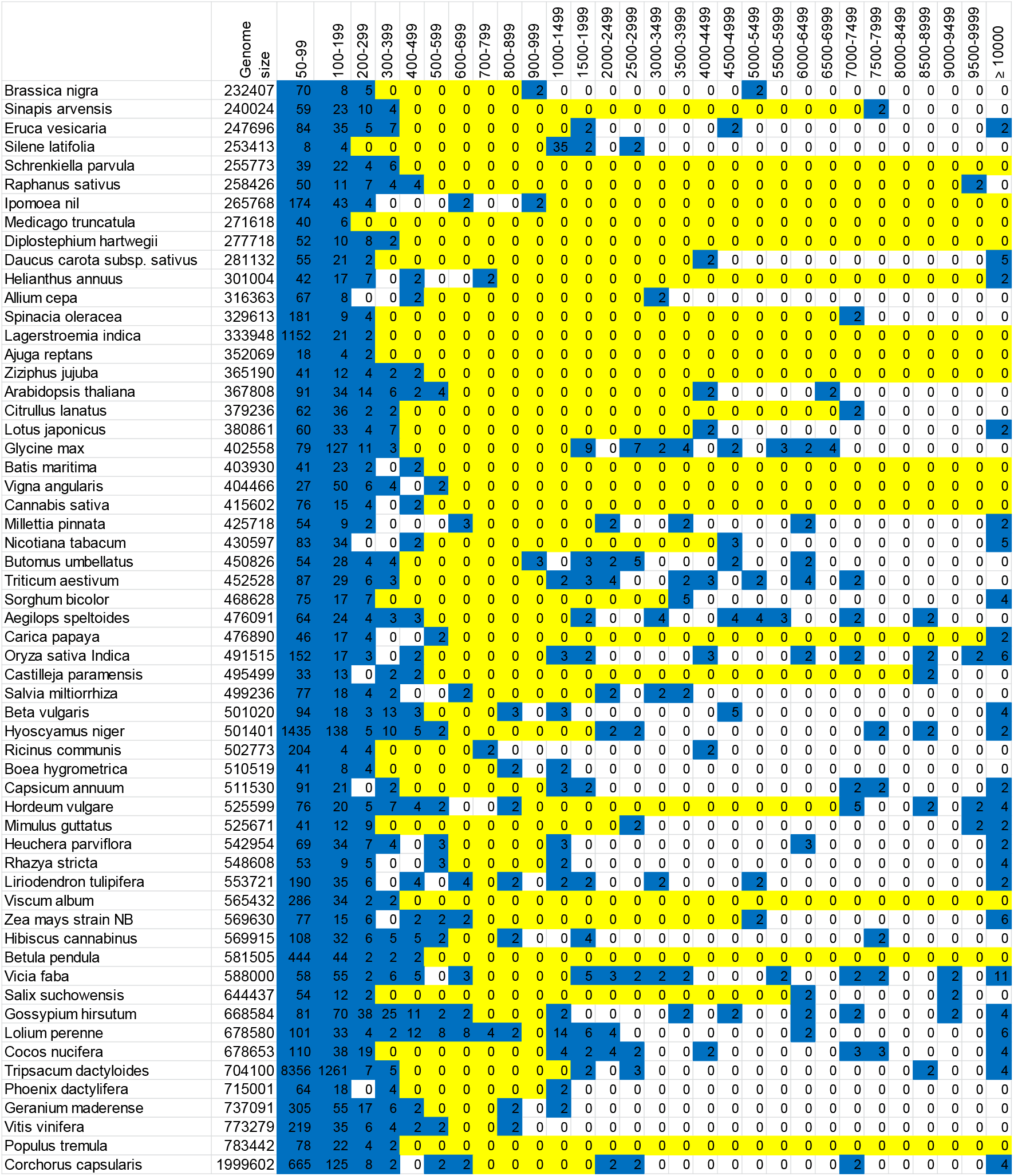
Distribution of repeat sizes among angiosperms. Species are sorted by increasing genome size. The number of repeats of each size class is shown. Blue shading indicates a number greater than zero. Yellow indicates a group of contiguous size ranges that contains zero repeats.

All eight of these species include one pair of long repeats, ranging in length from 1.9 to 9.7kb. However, these species fall into two groups. Each group has a large repeat that is found as a single copy in the other group. The B. nigra group consists of *R. sativus, S. arvensis, B. nigra* and *B. carinata;* the B. rapa group consists of *B. rapa, B. oleracea, B. napus* and *B. juncea*. A tree showing the relationships of these species, but without measures of distance, is shown in Figure 3. Part A shows the long repeat and neighboring sequences from the B. nigra group and the homologous single-copy sequences from the B. rapa group. Part B compares the long repeat from the B. rapa group to the unique homologous region from the B. nigra group.

**Figure 3.**
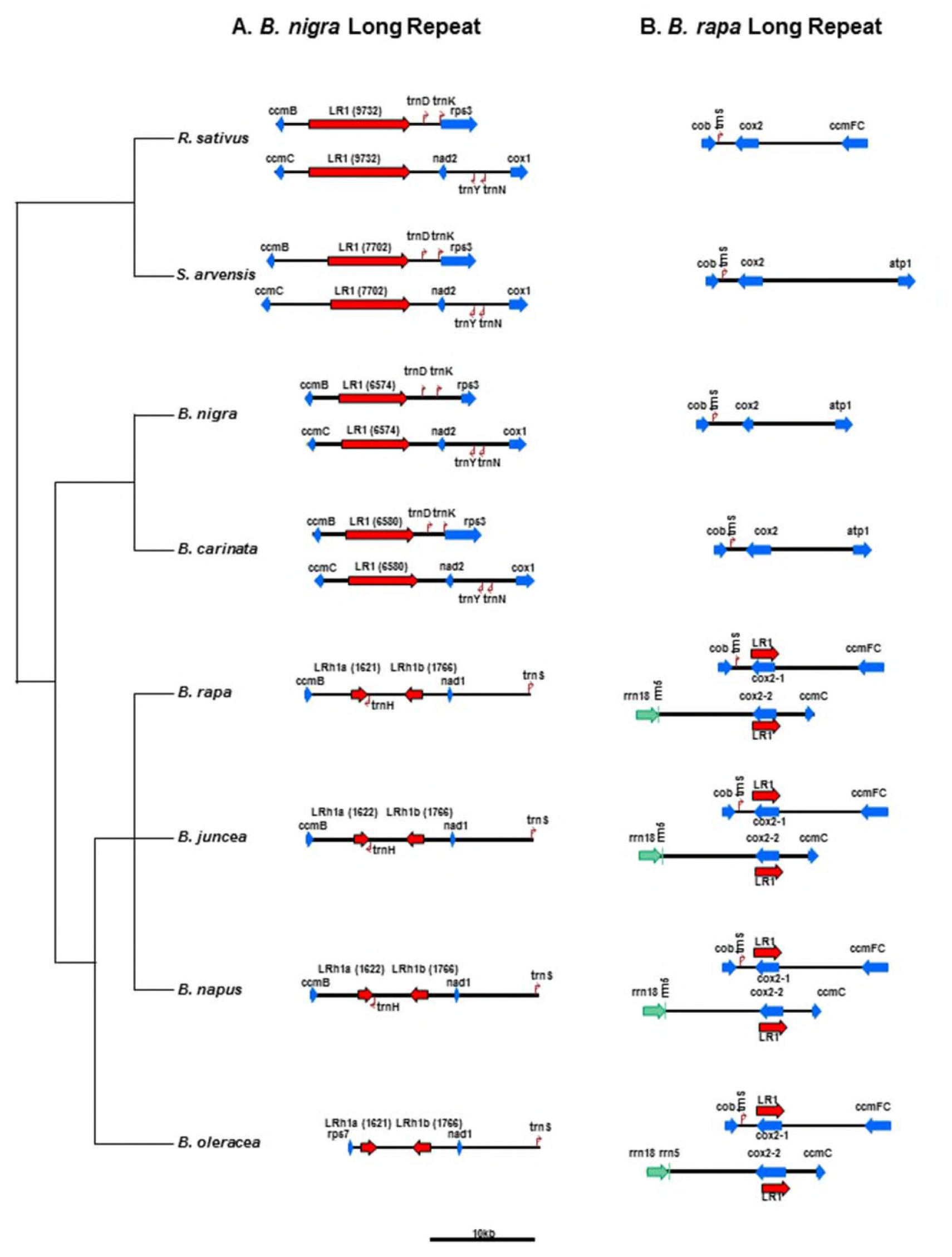
Long repeats in the Brassicales. A phylogenetic tree is shown at left, derived from Grewe *et al* (Grewe, F. *et al*. 2014). In part A are the regions surrounding the long repeat in *R. sativus, S. arvensis, B. nigra* and *B. carinata*. The homologus sequence from *B. rapa, B. napus, B. juncea* and *B. oleracea* is also shown. Part A shows the regions surrounding the long repeat in *B. rapa, B. napus, B. juncea* and *B. oleracea*, and the homologous region in the prior four species. Branch lengths in the tree are not to scale. The sequences are depicted at the scale shown in the figure.

The number of changes in coding DNA between plant species is generally low, making mutation rate estimates difficult, and these low rates may also be affected by sequencing errors (Sloan, D. B. *et al*. 2018). Grewe et al examined the synonymous substitution rates in genes of Brassicales mitochondrial genomes (Grewe, F. *et al*. 2014) and found them to be very low, consistent with most land plants. However, the presence of repeats allows mutations in non-coding DNA to be examined qualitatively. The long repeats of *R. sativus, S. arvensis, B. nigra* and *B. carinata* differ by large block substitutions and insertion/deletions (alignments are shown in Supplemental Figure S1). Where two copies are present in a species there are very few difference between copies, and they are generally near the boundaries of the repeats. Although significant differences can arise during speciation events, both copies of a repeat within a species remain identical. This supports the hypothesis that copies of repeated DNA are maintained as identical sequence by frequent recombination and gene conversion.

The long repeat of *B. nigra* and *B. carinata* underwent massive change in the lineage leading to the other four Brassica species. The first 1.6kb and the last 1.7kb of the repeat of *B. nigra* are conserved in the *B. rapa* group, and the *ccmB* gene still flanks the repeat on one side. However, the last 1.7kb are inverted and separated from the first 1.6kb by 3.3kb of a sequence of unknown origin. An additional difference is seen in *B. oleracea* wherein *rps7* now flanks the repeat rather than *ccmB*. Other major changes appear to have occurred in the time since *B. nigra* diverged from the ancestor of *B. oleracea* and *B. rapa;* a comparison of the complete mitochondrial genomes of *B. rapa* and *B. nigra* reveal at least 13 segments of DNA that have been rearranged. No major rearrangements have occurred between *B. nigra* and *B. carinata*, nor between *B. rapa, B. juncea* and *B. napus polima. B. oleracea* differs from *B. rapa* by approximately six rearrangement events.

At the same time that the *B. nigra* long repeat was being dramatically altered in the lineage leading to *B. rapa* and *B. oleracea*, a new long repeat appeared, which includes the coding sequence of the *cox2* gene. This new long repeat is maintained throughout this group of four species, and the flanking genes are also conserved (alignments are shown in Supplemental Figure S2). The *cox2* gene is single copy in *R sativus, S. arvensis, B. nigra* and *B. carinata*, and is in a nearly syntenic arrangement with neighboring genes.

## Discussion

The availability of complete mitochondrial genome sequences from many taxa of green plant allows us to compare the repeat structures across taxa. Although Large Repeats and ROUS have been known for some time, their functions (if any) and evolution are largely mysterious. It has been suggested that their existence and maintenance are outgrowths of double-strand break repair events such as nonhomologous end-joining (NHEJ), break-induced replication (BIR) and gene conversion (Christensen, A. C. 2018). We describe here a Python script that uses BLAST (Altschul, S. F. *et al*. 1990) to find non-tandem repeats within genomes, and use it to analyze plant mitochondrial DNA. Comparison of repeats between closely related species showed that repeat differences between species were largely due to rearrangements and block substitutions or insertions, which could be due to NHEJ and BIR, while the two copies of the repeat were identical within a species, suggesting continuing repair by gene conversion or homologous recombination.

The phylogenetic distribution of complex repeated structures in mitochondria appear to be common to the vascular plants and significantly different from the more primitive non-vascular taxa. This suggests that the common ancestor of lycophytes, ferns and seed plants adopted a new mechanism or strategy of mitochondrial genome replication and repair that led to a proliferation of repeats and increases in size. Complete sequences of more species, particularly in the lycophytes and ferns, is necessary to add clarity but the ancestor of vascular plants evidently made a transition to increased use of double-strand break repair in their mitochondria, leading to the genomic gymnastics seen today in plants.

The analysis of repeats in the *Brassica* species suggests that mitochondrial genomes can remain relatively static for long periods of time, but can also diverge very rapidly resembling punctuated equilibrium (Gould, S. J. and N. Eldredge 1977) that includes major rearrangements, sequence loss, and gain of sequences of unknown origin. The mechanisms and frequency are unknown, but it suggests that a lineage can experience a burst of genome recombination, breakage and rejoining, dramatically rearranging and altering the mitochondrial genome, as if it had been shattered and rebuilt. These events occur on a time scale that is faster than that of speciation, leading to the high levels of divergence, and a lack of strong correlations with the phylogeny.

Qualitative differences have been described between the repeats shorter and longer than about 1kb (Arrieta-Montiel, M. P. *et al*. 2009; Klein, M. *et al*. 1994; Mower, J. P. *et al*. 2012). The clustering within phylogenetic groups (Figure 2) and the trend towards bimodality (Figure 3) suggest differences between the large repeats and the smaller ones. We suggest that the term “large repeats” be reserved for those ROUS that are involved in genome isomerization. A working definition could be those ROUS larger than 1000bp, but functional analysis may reveal different size cutoffs in different species. Functional analysis can be done by analyzing clones big enough to include the repeats (Klein, M. *et al*. 1994), by long read sequencing (Shearman, J. R. *et al*. 2016) or Southern blotting. Functional analysis of the large repeats is an important step in understanding genome structure and evolution (Guo, W. *et al*. 2016; Guo, W. *et al*. 2017; Sloan, D. B. 2013) and may reveal different size cutoffs between species, which would reveal important differences in the replication and repair machinery and dynamics.

We doubt that there is an adaptive advantage to large size and abundant rearrangements in the genomes of plant mitochondria. We suggest that these are merely correlated traits accompanying the adaptive advantage of a greatly increased reliance on double-strand break repair. DNA repair is critically important because damage is more likely in mitochondria than the nucleus due to the changes in pH and redox potential, and the presence of reactive oxygen species. The strategy followed by animals is to minimize mutational targets by reducing genome size (Lynch, M. *et al*. 2006; Smith, D. R. 2016). However, with multiple copies of mitochondrial DNA in each cell, an alternative trajectory is to increase the use of template DNA in repair. The template-based accuracy of double-strand break repair is accompanied by the creation of chimeras, rearrangements and duplications when templates are not identical or cannot be found by the repair enzymes. Dramatic expansions, rearrangements and losses, accompanied by low substitution rates in genes is characteristic of flowering plant mitochondria. Selection on gene function maintains the genes, while the expansions and rearrangements must be nearly neutral. Once mitochondria evolved very efficient double-strand break repair, and a mechanism for inducing double-strand breaks at the sites of many types of damage, more primitive mechanisms, such as nucleotide excision repair can and have been lost (Gualberto, J. M. *et al*. 2014; Gualberto, J. M. and K. J. Newton 2017) without obvious evolutionary cost.

The adaptive value of increased and efficient double-strand break repair is probably to avoid mutations in the essential genes of mitochondria, and is possible because of the abundance of double-stranded template molecules in each cell. However this mechanism of repair has an additional correlated trait. There are bacterial species, such as *Deinococcus radiodurans*, that excel at double-strand break repair and can rebuild even significantly fragmented genomes (Krisko, A. and M. Radman 2013). This species is notoriously resistant to ionizing radiation, but the adaptive value of the trait is thought to be desiccation resistance, because dehydration also produces double-strand breaks (Mattimore, V. and J. R. Battista 1996). Radiation resistant bacteria in unrelated phylogenetic groups show more genome rearrangements and loss of synteny than their radiation sensitive relatives (Repar, J. *et al*. 2017), suggesting that abundant double-strand break repair is the cause of both the resistance to significant double-strand breakage and the loss of synteny. An interesting possibility is that very efficient double-strand break repair in plant mitochondria also confers desiccation resistance as a correlated trait. Because mitochondria are metabolically active immediately upon imbibition of seeds, DNA damage must be repaired very efficiently and rapidly (Paszkiewicz, G. *et al*. 2017). Efficient repair of desiccation-mediated damage in all cellular compartments is a prerequisite to being able to produce seeds or spores for reproduction. It is possible that the DNA repair strategy of plant mitochondria was one of several factors (including desiccation resistance of the nuclear and plastid genomes, presumably by distinct mechanisms) that are beneficial to vascular plants. The evidence of the repeats suggests that the transition to double-strand break repair in mitochondria occurred at approximately the same time as the transition to vascularity in plants, and it may have been one of several traits that enabled their success. In addition, once the life cycles of land plants included periods of desiccation in spores and seeds, double-strand breakage would have increased, accompanied by increases in rearrangements, expansions, and chimeras. The mechanisms of double-strand break repair continue to be important for understanding the evolution of plant mitochondrial genomes.

## Acknowledgements

We are grateful to Jeff Mower and Brandi Sigmon for many helpful comments on the manuscript, and Alex Kozik (UC Davis) for beta testing. ACC is grateful to Meric Lieberman and Isabelle Henry (U.C. Davis) for introducing him to Python scripting. ELW thanks Maya Khasin for support and encouragement. This work was supported in part by a grant from the National Science Foundation (MCB-1413152).

## Supplemental Materials

Supplemental File S1. Python script ROUSFinder.py

Supplemental File S2. Python script MultipleRepeats.py

Supplemental Table S1. Repeat sizes of all species used in this study. Bins include repeats larger than the size in the header, up to the next bin size.

Supplemental Figure S1. Alignment of the long repeats from *Raphanus sativus, Sinapis arvensis, Brassica nigra* and *Brassica carinata* with the homologous sequences from *Brassica rapa, Brassica juncea, Brassica napus polima* and *Brassica oleracea*. Part a is interleaved, and part b is in sequential fasta format.

Supplemental Figure S2. Alignment of the long repeats from *Brassica rapa, Brassica juncea, Brassica napus polima* and *Brassica oleracea*, with the homologous sequences from *Raphanus sativus, Sinapis arvensis, Brassica nigra* and *Brassica carinata*. Part a is interleaved, and part b is in sequential fasta format.

